# Traumatic brain injury disrupts state-dependent functional cortical connectivity in a mouse model

**DOI:** 10.1101/2023.10.06.560776

**Authors:** Samantha Bottom-Tanzer, Sofia Corella, Jochen Meyer, Mary Sommer, Luis Bolaños, Timothy Murphy, Sadi Quiñones, Shane Heiney, Matthew Shtrahman, Michael Whalen, Rachel Oren, Michael J. Higley, Jessica A. Cardin, Farzad Noubary, Moritz Armbruster, Chris Dulla

**Affiliations:** Department of Neuroscience, Tufts University School of Medicine, Boston, MA, USA; MD/PhD Program, Tufts University School of Medicine, Boston, MA, USA; Neuroscience Program, Tufts Graduate School of Biomedical Sciences, Boston, Massachusetts, USA; Department of Pathology, Case Western Reserve University School of Medicine, Cleveland, OH, USA; MD/PhD Program, Case Western Reserve University School of Medicine, Cleveland, OH, USA; Department of Neurology, Baylor College of Medicine, Houston, TX, USA; Department of Psychiatry, University of British Columbia, Vancouver, British Columbia, Canada; Djavad Mowafaghian Centre for Brain Health, University of British Columbia, Vancouver, British Columbia, Canada; Iowa Neuroscience Institute, University of Iowa, Iowa City, IA, USA; Department of Neurosciences, University of California San Diego, La Jolla, CA, USA; Department of Pediatrics, Harvard Medical School, Massachusetts General Hospital, Boston, MA, USA; Department of Neuroscience, Kavli Institute for Neuroscience, Yale School of Medicine, New Haven, CT, USA; Interdepartmental Neuroscience Program, Yale School of Medicine, New Haven, CT, USA; Department of Health Sciences, Northeastern University, Boston, MA, USA

**Keywords:** traumatic brain injury, controlled cortical impact, mesoscale imaging, in vivo GCaMP imaging

## Abstract

Traumatic brain injury (TBI) is the leading cause of death in young people and can cause cognitive and motor dysfunction and disruptions in functional connectivity between brain regions. In human TBI patients and rodent models of TBI, functional connectivity is decreased after injury. Recovery of connectivity after TBI is associated with improved cognition and memory, suggesting an important link between connectivity and functional outcome. We examined widespread alterations in functional connectivity following TBI using simultaneous widefield mesoscale GCaMP7c calcium imaging and electrocorticography (ECoG) in mice injured using the controlled cortical impact (CCI) model of TBI. Combining CCI with widefield cortical imaging provides us with unprecedented access to characterize network connectivity changes throughout the entire injured cortex over time. Our data demonstrate that CCI profoundly disrupts functional connectivity immediately after injury, followed by partial recovery over 3 weeks. Examining discrete periods of locomotion and stillness reveals that CCI alters functional connectivity and reduces theta power only during periods of behavioral stillness. Together, these findings demonstrate that TBI causes dynamic, behavioral state-dependent changes in functional connectivity and ECoG activity across the cortex.

## INTRODUCTION

Traumatic brain injury (TBI) is the leading cause of death and disability among trauma-related injuries.^1^ Following moderate to severe TBI, there is significant mortality and less than a quarter of patients experience functional recovery in a 5-year post-injury window.^2^ TBI causes cognitive and motor dysfunction,^3–5^ can lead to post-traumatic epilepsy (PTE),^6–8^ and increases risk for neurodegenerative diseases,^9,10^ all conditions associated with network dysfunction.^11,12^ Although multiple molecular and cellular changes are known to occur after TBI, the functional impact of TBI on neuronal activity and circuit function remains poorly understood. Elucidating how the brain responds to injury on a network level is thus a key goal for identifying biomarkers of adverse outcomes and developing targeted therapeutic interventions.

Functional connectivity is a measure describing the short-latency temporal correlation of neuronal activity between brain regions and can provide information about how networks behave at baseline, during task performance, and after different pathophysiologic perturbations including TBI.^13–18^ Functional connectivity can be measured using electroencephalography (EEG), electrocorticography (ECoG), magnetoencephalography (MEG), magnetic resonance imaging (MRI), and mesoscale imaging.^11,13–15,19–24^ MEG and fMRI analysis of contusional TBI patients reveals decreased functional connectivity in damaged tissue compared to contralateral uninjured tissue and to controls.^14,15,25^ Recovery of functional connectivity after TBI is associated with improved cognition and memory both globally and within networks responsible for measurable cognitive functions.^11,15,26^ Functional connectivity patterns of TBI patients who underwent rehabilitation were more similar to healthy controls than to non-rehabilitated TBI patients and showed better functional recovery.^15^

To assay functional connectivity in a clinically relevant mouse model of TBI, we developed a novel approach to perform in vivo widefield cortical mesoscale imaging with simultaneous ECoG in mice after controlled cortical impact (CCI). CCI is an established rodent model of TBI that primarily models contusion and hemorrhage, and leads to circuit reorganization, in vitro hyperexcitability, and post-traumatic epilepsy with a similar incidence to what occurs in patients with comparable injury severity.^6,12,27–33^ Among patients with severe TBI, up to 35% experience intracerebral contusion and are the most neurologically impaired, have the highest healthcare impact, and the greatest societal impact.^34,35^ Mesoscale imaging allows in vivo widefield imaging of the cortical surface in awake, behaving animals.^21,22,36,37^ We quantified neuronal activity by using neuronal expression of the fluorescent reporter GCaMP7c, a genetically encoded calcium indicator.^38^

Not only is functional connectivity altered with injury, but it is highly dynamic across brain states. When at rest, cortical functional connectivity is highest, and it is reduced during increases in attention and motor-demanding tasks.^19,39,40^ Measuring functional connectivity during different brain states enables a more complete understanding of how discrete brain regions are activated, recruited, and coordinated across states. However, little is known about how functional connectivity changes as a function of brain state in the injured brain. Therefore, we carried out mesoscale imaging while mice were free to run on a wheel and tracked their behavioral state (locomotion versus stillness) to understand state-dependent functional connectivity in the injured brain.

We report that CCI disrupts coherent cortical activity most profoundly immediately after injury in regions proximal to the lesion including somatosensory, parietal, and primary visual cortices. This is followed by a partial recovery during the weeks that follow. We also identify state-dependent changes in both ECoG power and functional activity during locomotion versus stillness that are disturbed by CCI. Interestingly, the biggest effects of CCI on functional connectivity are seen during stillness, with much smaller changes in functional connectivity during movement. Together, we report a novel approach to carry out mesoscale imaging with simultaneous ECoG after CCI and show temporally, regionally, and state-specific changes in functional connectivity in the injured rodent brain.

## METHODS

### Animals

Male CD-1 mice (bred in house from Charles River Labs stock, strain code 022) were kept on a 12-hour light/dark cycle with access to food and water ad libitum. Following surgeries, mice were singly housed. All procedures were approved by the Tufts University Institutional Animal Care and Use Committee.

### Focal viral injections

Cortical expression of the calcium sensor GCaMP7c was attained via focal viral injection of AAV1 under the human synapsin (hsyn) promoter for broad neuronal expression (#105321, Addgene).^41^ CD-1 male mice (P42-56) were stereotactically injected in both hemispheres using three injection sites/hemisphere (coordinates): (+/-2, 1.5, 0.75), (+/-2, 3.5, 0.75), and (+/-1.75, 6, 0.75) (λ + x, +y, −z) mm. Mice were anesthetized with isoflurane (4% induction, 2% maintenance) and body temperature was maintained on a heating pad at 37⁰C. Buprenorphine (0.1 mg/kg) was injected subcutaneously. A small incision was made, and burr holes drilled. ∼5×10^10^ viral gene copies were injected per site (1 μl per site, 1:1 dilution with saline). The skin incision was closed using surgical sutures (#359551, McKesson). Mice were monitored closely until they resumed normal behavior and then were monitored daily for three days to ensure recovery. Mice were used for mesoscale implant 14-21 days following injection.

### Surgical procedures

For implantation of mesoscale window and ECoG, anesthesia was induced, body temperature maintained, and analgesic administered as above. The skin and fascia overlying the skull from the interparietal bone to the nasal bone and between the temporal muscles was removed. The skull was thinned using a 0.7 mm spherical burr (FST) and dental drill. The skull was appropriately thin when pliable under curved forceps and vasculature could be easily visualized. Next, mice were randomly selected to receive either CCI or sham injury. For CCI, a 5 mm craniectomy was made over the left somatosensory cortex using a dental drill and trephine. CCI was delivered using a Leica Benchmark Stereotaxic Impactor (3 mm piston, 1 mm depth, 400 ms dwell time, 3.5 m/s velocity). The craniectomy was sealed with cyanoacrylate (3M). Sham mice received skull thinning but no craniectomy or CCI. Clear dental cement (C&B Metabond, Parkell) was applied, and a custom cut glass window covering the craniectomy and entire thinned skull was implanted (diamond scriber, VWR; No 1 gold seal cover glass, Thermo Scientific). Burr holes were drilled and ECoG electrodes implanted (#8403, Pinnacle) bilaterally over the somatosensory cortex laterally to the window (coordinates): (+/-4.5, 4) (λ + x, +y) mm. Ground and reference electrodes were implanted in intraparietal bone. ECoG electrodes were soldered to a headmount (#8402, Pinnacle). Cold cure dental cement (Teets Denture Material, Cooralite Dental Manufacturing) mixed 2:1 with black tempera paint powder (Jack Richeson & Co.) was used to secure ECoG electrodes, custom head bar, and the glass window and to decrease optical interference. The head bar was anchored posterior to the window over ground and reference electrodes. Animals recovered on a warming pad until they resumed normal behavior and then were monitored daily to ensure post-surgical recovery.

### Immunohistochemistry

Mice were deeply anesthetized and transcardially perfused with PBS followed by 4% paraformaldehyde (PFA) in 0.4M phosphate buffer. Brains were extracted, post-fixed in PFA for 24 hours, and then cryopreserved in 30% sucrose. Coronal slices (40 μm) were made on a Thermo Fisher Scientific Microm HM 525 cryostat. Free-floating sections were blocked in 10% goat serum (GS), 5% bovine serum albumin (BSA), and 0.2% TWEEN20 -PBS (PBST) at room temperature for 1 hour. Sections were immunolabeled with primary antibodies in 5% GS/1% BSA/0.2% PBST. Primary antibodies were used as follows: Iba-1 (1:1000; Wako 019-19741), GFAP (1:500; IF03L-100UG), GFP (1:1000; Abcam ab13970). Secondary antibodies were used at a ratio of 1:500 in 5% GS/1% BSA/0.2% PBST (anti-rabbit-Alexa-Fluor-555, Invitrogen A27039; anti-mouse-Alexa-Fluor-647, Invitrogen A21235; anti-chicken-Alexa-Fluor, Abcam ab150169). Images were collected for analysis using a Keyence epifluorescence microscope with a 10X objective from a single plane of focus per section, and composite images of hemi-cortex were made. Illumination intensity and exposure time were kept constant across all image collection. For image analysis, regions of interest (ROIs) were drawn using Image J software (NIH) in primary somatosensory cortex. Three sections per animal with one ROI per section were quantified. For CCI-injured mice, ROIs were drawn on the cortex ipsilateral to injury from the midline to beyond the lateral edge of the cavitary lesion superficial to CA3. ROIs were drawn in identical cortical regions in shams. Quantification of Iba-1+ cell density within each ROI and fluorescence intensity of Iba-1+ cells were performed using CellProfiler. Global GFAP expression within each ROI was quantified using a custom MATLAB script to determine the percent area that was GFAP positive.^23^ GFP was quantified using the mean fluorescence intensity within ROIs. Representative images for Figure 3 were collected using a Nikon A1R confocal microscope with a 40X oil objective (Olympus).

### Widefield imaging and ECoG

Widefield calcium imaging was performed using a high-speed high-resolution Andor Zyla sCMOS camera with 470nm and 535nm alternating excitation (mounted LEDs, ThorLabs) to image GCaMP7c and hemodynamic reflectance, respectively.^21,42^ Emitted light from the 470nm LED passed through a 460/50nm excitation filter (ET460/50m, Chroma) and from the 530nm LED through an absorptive filter (NE10A, ThorLabs). Light then passed through (from 530nm source) or was reflected off (from 470nm source) a 495nm dichroic mirror (T495lpxr, Chroma), a 470/530nm dual excitation filter (CT470/530x, Chroma), and a 525/36nm emission filter (ET525/36m, Chroma). Images were acquired at 256 x 256 resolution, and each channel was acquired at 33 Hz with 5 ms exposure time. ECoG recording was performed using a PowerLab 8/35 (ADInstruments) at a sampling rate of 1000 Hz recorded in LabChart 8 (ADInstruments). Mice were head-fixed on a custom cylindrical running wheel and allowed to acclimate for 30 minutes prior to each imaging session.^43,44^ Mice did not undergo head fixation prior to 3 day post-injury (DPI) to allow for post-surgical healing. A magnetic encoder (MAE3, US Digital) was used to record rotational movement of the wheel. ECoG was recorded throughout the entire session and images were collected over 30 minutes (6 x 75 second imaging periods). Sessions were performed during the light phase of the 12-hour light/dark cycle at 3, 5, 7, 14, and 21 DPI.

### Mesoscale data analysis

#### Preprocessing of imaging and ECoG data

All analyses were conducted using custom-written scripts in MATLAB version 2022b. ECoG data was imported into MATLAB from LabChart 8 and included ECoG from two channels, locomotion data, and timing of alternating LED triggering. Imaging trials were identified based on LED triggering (66 Hz for 75 seconds) and aligned with ECoG. Locomotion was identified from stillness by setting a minimum value for the integrated voltage change over 100 ms as detected by the running wheel encoder. Images were separated into fluorescence (470nm excitation) and reflectance (530nm excitation). Images were spatially aligned to bregma and the cross point between the median line connecting the frontal poles. ^45^ A mask was used to define cortical boundaries and applied to the full imaging stack.

#### Normalization and hemodynamic correction

To normalize fluorescence (F) and reflectance (R) data, ΔF/F _0_ and ΔR/R_0_ for each pixel was calculated using (F - F_0_)/F_0_, where F is the raw unfiltered signal and F _0_ is the mean of the full image stack.^21,46^ This was repeated for reflectance. To correct for hemodynamic artifacts, normalized reflectance was subtracted from normalized fluorescence as ΔF/F_0_ - ΔR/R_0_. ΔF/F_0_ - ΔR/R_0_ values were used for all subsequent analyses.^21^

#### Postprocessing analysis of imaging data

For anatomic mapping, images were aligned and anatomic landmarks registered as above. Ten cortical ROIs (5/hemisphere) were mapped onto images using Paxinos and Franklin’s stereotactic coordinates including: secondary motor cortex, motor barrel cortex, primary somatosensory area forelimb, parietal cortex, primary visual cortex. The ROIs were small (radius = 3 pixels) ensuring they were entirely within the indicated anatomic regions. For assessment of total neuronal activity, a trace from each ROI was generated representing fluorescent activity over time and the coefficient of variation (CV) of each trace calculated. This was repeated for all imaging sessions to generate a mean CV value for each brain region at every timepoint. Pearson’s correlation coefficients between ΔF/F_0_ - ΔR/R_0_ signals were computed using the “corrcoef” function (MATLAB) which is amplitude independent.^45^ Correlation coefficients were displayed as connectivity matrices. This process was repeated for all imaging trials and all behaviorally-defined segments at least 8 seconds in length of locomotion and stillness within each imaging trial. To eliminate effects of state transition, we removed the first 1.5 seconds and last 0.5 seconds of each period of stillness or locomotion.^22^ We measured interhemispheric connectivity by computing the correlation coefficient between left primary motor cortex and all 5 contralateral regions. This was repeated for the 9 remaining regions and the mean across all regions was taken. For intrahemispheric connectivity, the correlation coefficient between regions within each hemisphere was computed and the mean taken.

#### Postprocessing analysis of ECoG data

To quantify ECoG power, raw ECoG signal was bandpass filtered using the “bandpower” function (MATLAB) into five bands: total (1 – 100 Hz), low (2 – 6 Hz), theta (6 – 8.3 Hz), alpha (8 – 12 Hz), and gamma (30 – 70 Hz).^47^ Bands were normalized to total power. Power from these bands was identified from the entire ECoG recording, the first half, and the second half to determine any effect due to acclimation to the wheel, and each imaging trial.

### Statistics and reproducibility

Statistical analyses were conducted using custom-written scripts in MATLAB and RStudio (2022.07.2 Build 576). Figures were created in Prism 8 (GraphPad) and on BioRender.com. Three animals were used in each treatment group. For all experiments, we performed linear mixed modeling (LMM) to measure statistical differences due to animal injury status, state, and DPI (fixed effects) while accounting for intra- and inter-animal variability (random effects).^31,48^ Dependent variables included GFAP fluorescence, Iba1 density (Figure 2), ΔF/F_0_ (Figure 4), connectivity (Figures 5, 7), and ECoG power (Figure 6). The “lmer” function (RStudio) was used for running LMM with lme4, lmerTest, matrix, and stats packages. For LMM, t values >1.96 and <-1.96 were considered statistically significant. Example code is as follows: corrcoef = lmer(corr_coef ∼ injury_status * dpi + (1|trial_num:mouse_num) + (1|mouse_num), data = Connect_Interhem, na.action = “na.omit”). In this example from Figure 5, we tested the data (Connect_Interhem) for an effect of injury (injury_status) and time post-injury (dpi) on functional connectivity (corr_coef). Inter-animal (1|mouse_num) and intra-animal (1|trial_num:mouse_num) variability were considered. In figures 5 and 7, average differences in connectivity matrices were determined using a Wilcoxon rank sum test (unpaired) or Wilcoxon signed rank test (paired) and multiple comparisons controlled for using a Benjamini–Yekutieli correction.^22^ No animals were excluded from analysis. Post hoc power analysis was performed using G*Power (3.1.9.7) on the data in Figure 5A to confirm that sample sizes are sufficient given the effect sizes observed. ^49,50^ Input parameters: effect size f = 30.246, α = 0.05, total sample size = 6, number of groups = 2, number of measurements = 18, corr among rep measures = 0.05. Output parameters: noncentrality parameter λ = 10400.06, critical F = 7.7, numerator df = 1, denominator df = 4, power = 1. Statistical results are reported in their entirety in Tables S1 and S2.

## RESULTS

### Simultaneous mesoscopic imaging and ECoG in CCI-injured mice

To perform widefield cortical imaging in the CCI model of TBI, we developed a novel surgical and imaging approach for simultaneous GCaMP mesoscale imaging and ECoG in awake mice following TBI. Our protocol allows mice to run freely on a running wheel to analyze state-specific changes in brain activity during stillness and motion. Focal cortical injections (three/hemisphere) of the neuronal calcium indicator GCaMP7c (AAV1-syn-jGCaMP7c), selected for its high contrast and low baseline fluorescence,^38^ were made in adult male CD-1 mice. This resulted in wide reporter expression throughout both cortical hemispheres including somatosensory, retrosplenial, motor, and visual cortices. Two weeks later, mice received sham or CCI injury and were implanted with a cortical window, ECoG electrodes, and a head bar for head fixation (Figure 1A). Our implant approach does not require any modification of CCI parameters and allows optical access to the dorsal surface of the cortex including the site of injury. Beginning 3 DPI to allow time for recovery after surgery, mice were head-fixed on a custom-made mesoscale imaging microscope with a running wheel and allowed to run freely during imaging and ECoG recording. Mice were imaged and ECoG recorded 3, 5, 7, 14 and 21 DPI.

**Figure 1.**
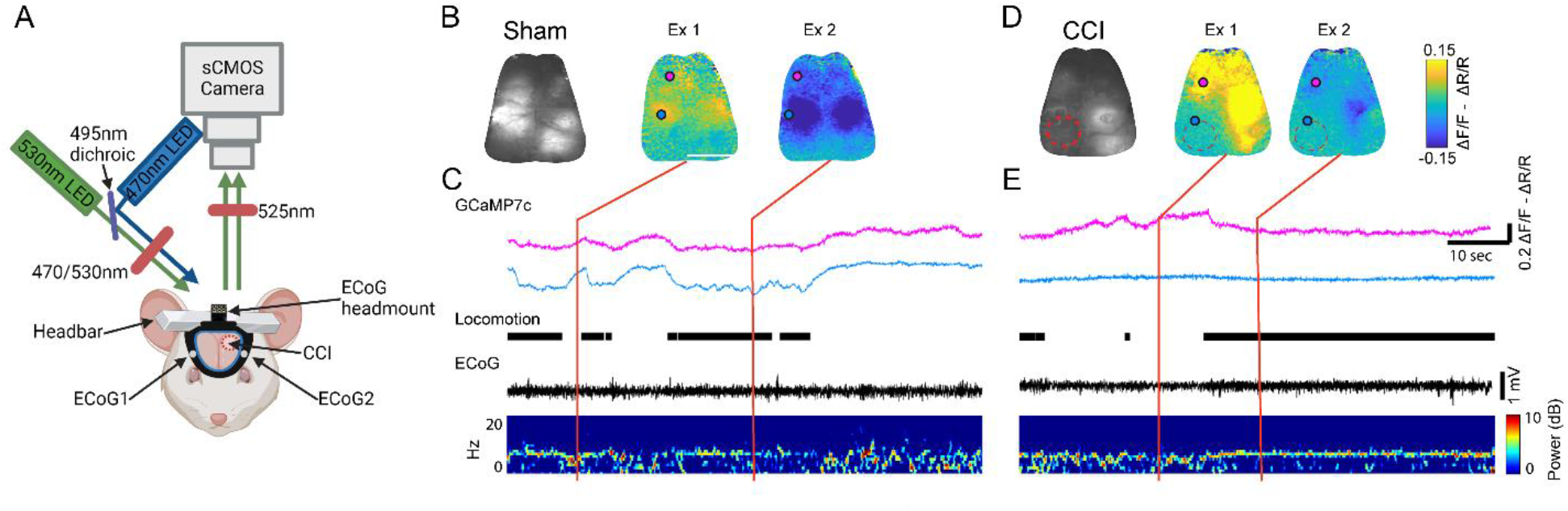
Implant design, imaging setup, and spatiotemporal cortical dynamics of neuronal activity and ECoG. **A.** Diagram of the implant & imaging setup including camera, LEDs, and ECoG recording equipment. **B.** Baseline image of sham implanted mouse (left) and example normalized images showing fluctuations in neuronal (GCaMP7c) activity during stillness (Ex 1) and during movement (Ex 2). Overlaid circles indicate anatomic regions from which activity is quantified in the traces below. Pink indicates secondary motor cortex (M2) and blue indicates primary somatosensory area, forelimb (SSC). Scale bar = 3mm. **C.** From top to bottom, sample traces of GCaMP7c activity from M2 (pink) and SSC (blue), times of locomotion (black bar) for the animal shown in B, ECoG, and corresponding spectrogram. Red lines indicate temporal alignment with Ex 1 and 2. **D.** Same as B but of a CCI-injured mouse. Red dotted circles indicate location of CCI. **E.** As in C but for the CCI animal indicated in D. These example images come from 21 days after CCI or sham surgery.

Images were anatomically registered using visually identifiable anchoring coordinates.^45,51^ Fluorescence and reflectance data were separated and normalized (Figure 1B, 1D).^52^ Reflectance was subtracted from fluorescence to account for hemodynamic fluctuations. ^21^ All analyses utilized this normalized and corrected signal to account for small differences in baseline fluorescence due to focal viral injections and curvature of the brain. ECoG signal was temporally aligned with imaging data and behavior state was categorized as locomotion or stillness (Figure 1C, 1E). Throughout the cortex in sham and injured animals, large, spatially specific calcium signals were seen, indicative of neuronal activity. Mice spontaneously transitioned between stillness and locomotion, and movement state corresponded to fluctuations in calcium activity and ECoG spectrogram. In ECoG traces, there was prominent theta activity during periods of locomotion. Thus, we demonstrate that the CCI model of TBI can be integrated with widefield calcium imaging and ECoG.

### Mesoscale implant slightly increases GFAP but not Iba1 immunolabeling after CCI

We next asked if implanting a mesoscale window altered the response to CCI, as quantified by changes in glial reactivity. Glial cells, including astrocytes and microglia, mount the primary response to brain insult and contribute to secondary injury effects following TBI.^53,54^ After injury, astrocytes can become reactive and cause inflammation, alter the blood-brain barrier, and form a glial scar.^55–57^ Microglia are activated, recruited to the site of injury, secrete inflammatory molecules, and are associated with post-injury cognitive deficits.^58–60^ To address whether mesoscale implant alters glial reactivity after CCI, we quantified the expression of markers for astrocyte (GFAP) and microglia (Iba1) reactivity.

Animals underwent CCI or sham injury, and half received subsequent mesoscale implant. Three weeks later, brains were prepared for immunolabeling with GFAP and Iba1 as a proxy for astrocytosis (as assayed by total area of GFAP immunohistochemical staining) and microglial activation (as measured by the number of Iba1+ cells and the intensity of Iba1 staining per cell), respectively (Figure 2A). We found a significant increase in GFAP expression and Iba1+ cell number in CCI mice, as compared to shams, in both implanted and un-implanted groups, consistent with abundant glial reactivity after TBI (GFAP: t_(8)_ = 2.39, p = 0.04 effect of implant; t_(8)_ = 5.19, p = 8e-4 effect of injury; t_(8)_ = 2.35, p = 0.04 interaction effect, LMM) (Iba1: t_(8)_ = 3.18, p = 0.01 effect of injury, LMM) (Figure 2D,E; Table S1).^57^ Mesoscale implant caused a small but significant increase in GFAP expression, but not number of Iba1+ cells, in CCI-injured mice alone. It is only in the context of a concurrent CCI that window implantation increases GFAP expression. Window implantation alone does not elicit an increase in GFAP.

**Figure 2.**
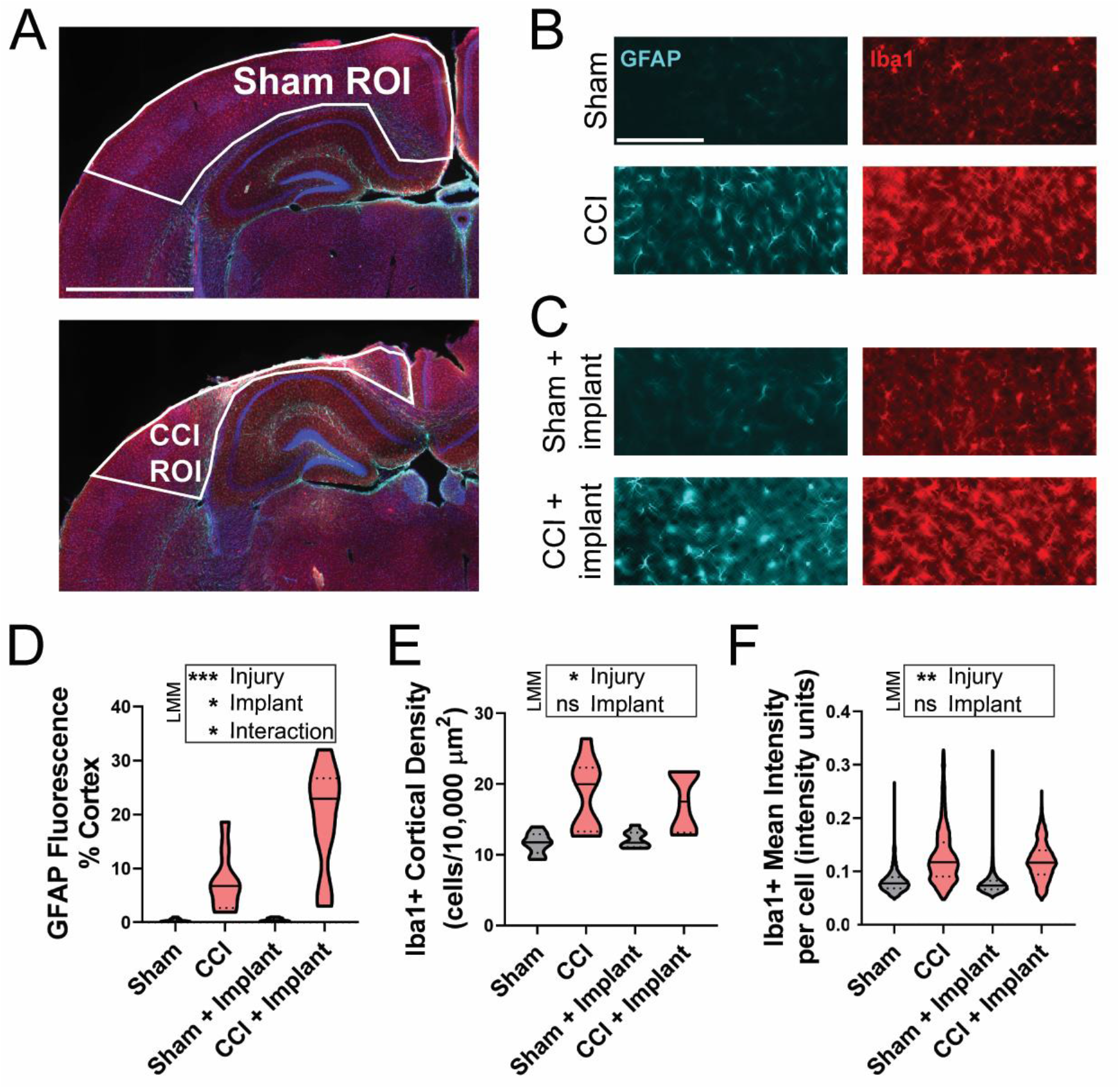
Mesoscale implant slightly increases GFAP but not Iba1 immunolabeling after CCI. **A.** Example ROIs drawn in cortex of a sham (top) and a CCI (bottom) mouse brain section that were used for image quantification in D-F. Scale bar = 1000 µm. **B.** Representative fluorescent images from somatosensory cortex of non-implanted sham and CCI mice 21 days post-surgery immunolabeled for GFAP (cyan) and Iba1 (red). For CCI mice, images are from cortex ipsilateral to injury. Scale bar = 100 µm. **C.** Representative fluorescent images of sham and CCI mice implanted with a cranial window 21 days post-surgery immunolabeled for GFAP (cyan) and Iba1 (red). **D.** Injury and mesoscale implant both significantly increased GFAP expression in cortex, quantified as area of cortex that was GFAP positive. **E.** Mean cortical density of Iba1+ cells were significantly increased by injury with no apparent effect of mesoscale implant. **F.** Mean Iba1+ cell fluorescence intensity was increased by injury, but not by mesoscale implant. Total number of cells are as follows: Sham: 20,064, CCI: 14,378, Sham + Implant: 18,700, CCI + Implant: 13,940. For all plots, n = 3 animals per group, 3 sections per animal. LMM: *indicates P < 0.05, **indicates P < 0.01, ***indicates P < 0.001. In violin plots, median is indicated with a solid line, quartiles are demarcated with dashed lines.

Similarly, when comparing the Iba1+ intensity within each cell, CCI but not implant caused a significant increase (t_(8)_ = 4.42, p = 2e-3 effect of injury, LMM; Figure 2F). This shows that CCI significantly increases GFAP and Iba1 immunoreactivity, consistent with reactive astrocytosis and microglial activation after injury. The addition of the mesoscale implant causes a small but significant increase in GFAP but not Iba1 immunolabeling in the injured cortex, indicating that mesoscale implant slightly increases astrocyte reactivity after CCI.

### GCaMP7c expression is not significantly altered by CCI

To test whether CCI altered the expression of GCaMP7c, we performed CCI and sham surgeries on animals focally injected with GCaMP7c and then used immunohistochemistry and epifluorescence imaging to quantify GCaMP7c expression independent of neuronal activity. 21 days post-surgery, brains were prepared for immunohistochemical analysis of reporter expression using an anti-GFP antibody. GFP immunoreactivity was quantified in cortical layers I-III corresponding to the area of cortex that was imaged during mesoscale imaging (Figure 3A,B).^36^ There were no significant differences in GFP immunoreactivity between CCI and sham mice (t_(4)_ = −1.09, p = 0.34, LMM). There were also no significant differences in GFP immunoreactivity in layers IV-VI (data not shown). We saw persistent GCaMP7c expression in injured animals adjacent to the site of injury three weeks after injury (Figure 3C). This supports that any changes in GCaMP7c activity proximal to injury are not due to decreased reporter expression, but rather occur due to changes in neuronal activity.

**Figure 3.**
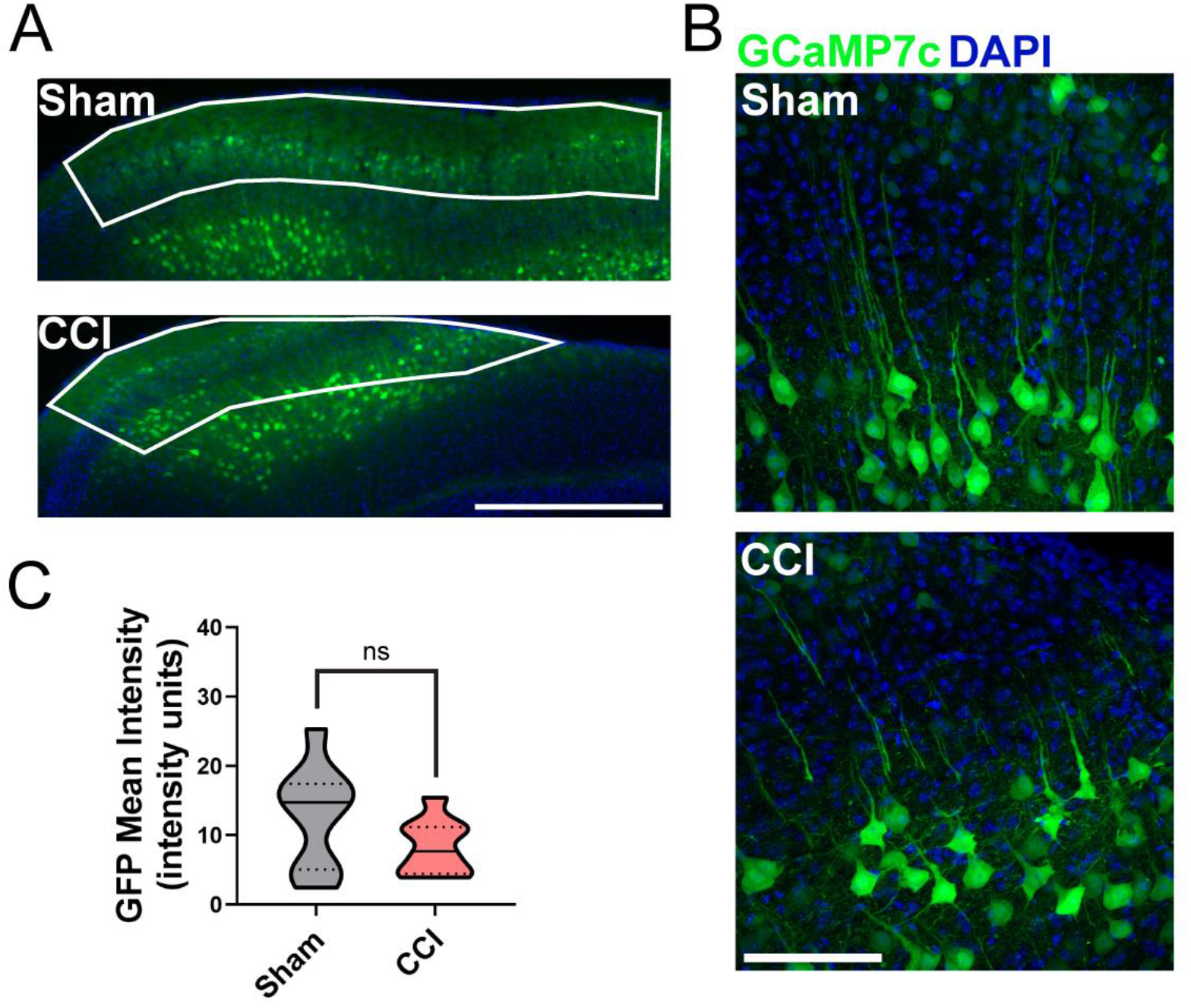
GCaMP7c expression is not significantly altered by CCI. **A.** Representative fluorescent images of sham (top) and CCI (bottom) injured mice 21 days post-surgery immunolabeled for GFP (green). Example ROIs drawn in the cortices were used for image quantification in B. Scale bar = 500 µm. **B.** Representative confocal images of GFP-expressing cells in layers I-III in a sham (top) and CCI (bottom) mouse. Note that the CCI representative image is perilesional. Scale bar = 100 µm. **C.** Mean GFP fluorescence intensity in layers I-III was not significantly affected by CCI. For plot, n = 3 animals per group, 3 sections per animal. LMM: ns is not significant. In violin plot, median is indicated with a solid line, quartiles are demarcated with dashed lines.

### Neuronal activity is decreased proximal to CCI and increased in the contralateral cortex

To determine whether CCI affected total neuronal activity during imaging, the coefficient of variation (CV) of GCaMP7c fluorescence was quantified in each imaging session across multiple cortical regions. CV allows a simple quantification that integrates both the dynamic range and the total ongoing activity during imaging.^61^ Region of interest (ROI) placement was automated and ROIs were placed bilaterally to include motor, somatosensory, parietal, and visual cortices. CCI border was marked ensuring ROIs proximal to CCI were perilesional, not in the injury itself (Figure 4A). A mean CV value for each brain region at every timepoint was generated (Figure 4B). Following CCI, GCaMP7c CV was significantly decreased over all timepoints examined in the three brain regions most proximal to CCI as compared to identical regions in sham-injured mice (t_(43)_ = −6.12, p = 2.5e-7, LMM; Figure 4C).

**Figure 4.**
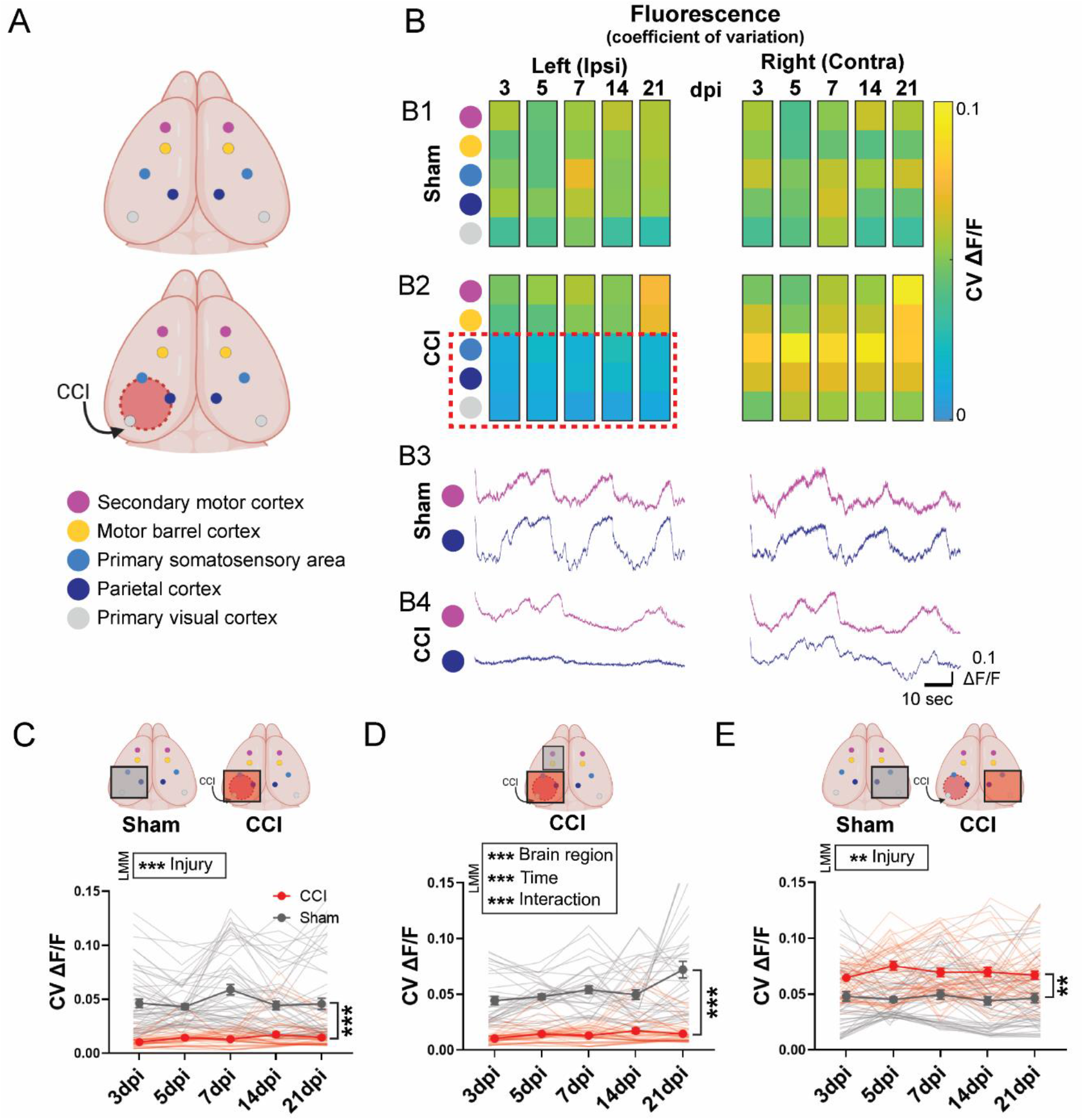
Neuronal activity is decreased proximal to CCI and increased in the contralateral cortex. **A.** Schematic of location of ROIs in sham (top) and CCI (bottom). Colors correspond with panels in B and are anterior to posterior: secondary motor cortex (pink), motor barrel cortex (yellow), primary somatosensory area forelimb (light blue), parietal cortex (dark blue), and primary visual cortex (gray). **B.** Coefficient of variation (CV) of ΔF/F_0_ in the regions mapped in A in sham (B1) and CCI (B2) mice. Red dashed box indicates perilesional regions. B3. Sample ΔF/F _0_ traces from a single imaging session 3dpi from sham and CCI (B4) mice. **C.** Mean CV of ΔF/F_0_ from the three ROIs proximal to CCI (demarcated with red box) compared to identical regions in sham mice (ROIs within grey box). Lines indicate all replicates contributing to the mean. **D.** Mean CV of ΔF/F_0_ from the three ROIs proximal to CCI (red box) compared to ROIs anterior to CCI (grey box). **E.** Mean CV of ΔF/F_0_ from ROIs contralateral to CCI (red box) compared to identical regions in sham mice (grey box). For C-E, LMM: *indicates P < 0.05, **indicates P < 0.01, ***indicates P < 0.001. Error bars indicate SEM. Dpi, days post-injury. Ipsi, ipsilateral. Contra, contralateral.

Interestingly, this decrease in GCaMP7c CV was limited to cortical areas just outside the CCI and was not seen in regions anterior to injury (t_(429)_ = 9.23, p < 2e-16 effect of brain region; t_(429)_ = 5.66, p = 2.84e-8 effect of time; t_(429)_ = 4.1, p = 5.9e-5 interaction effect, LMM; Figure 4D). Finally, regions contralateral to CCI had an increased GCaMP7c CV when compared to matching regions in sham-injured mice at all timepoints (t_(56)_ = 2.96, p = 4.5e-3 effect of injury, LMM; Figure 4E). Overall, GCaMP7c CV was consistently decreased in somatosensory, parietal, and primary visual cortices ipsilateral to injury, and increased in contralateral somatosensory, parietal, and primary visual cortices following CCI.

### CCI regionally impairs functional cortical connectivity that partially recovers over time

We next asked whether CCI alters functional connectivity, as measured by coherent GCaMP7c activity across brain regions. Interhemispheric connectivity, quantified using the strength of correlated neuronal activity across hemispheres, is an important readout of functional connectivity and is altered with TBI.^26^ We measured interhemispheric connectivity by mapping the same 10 ROIs onto the cortex as in Figure 4 and computed the correlation coefficient between all regions. We found that CCI significantly decreased interhemispheric functional connectivity (t_(15)_ = −6.64, p = 9.2e-6 effect of injury; t_(170)_ = 6.0, p = 1.18e-8 interaction effect, LMM; Figure 5A). This was evaluated 3, 5, 7, 14, and 21 days post-injury. To determine if changes in functional connectivity were driven by changes in the injured versus contralateral hemispheres, we also calculated intrahemispheric connectivity. We computed the average functional connectivity between all regions in the left hemisphere and separately for the right hemisphere (Figure 5B). Interestingly, CCI reduced functional connectivity only in the injured hemisphere (t_(5)_ = −6.22, p = 2.3e-3 effect of injury; t_(172)_ = 8.39, p = 1.7e-14 interaction effect, LMM). Connectivity in the injured hemisphere significantly recovered over time, more than doubling over the 21 days following injury. There was no significant decrease in connectivity in the hemisphere contralateral to injury after TBI, only a small effect of time indicating some variability in connectivity in the contralateral cortex over time (t_(172)_ = −2.43, p = 0.02 effect of time, LMM). Together, these data indicate that CCI drives largely focal disruptions in functional connectivity that recover over time.

**Figure 5.**
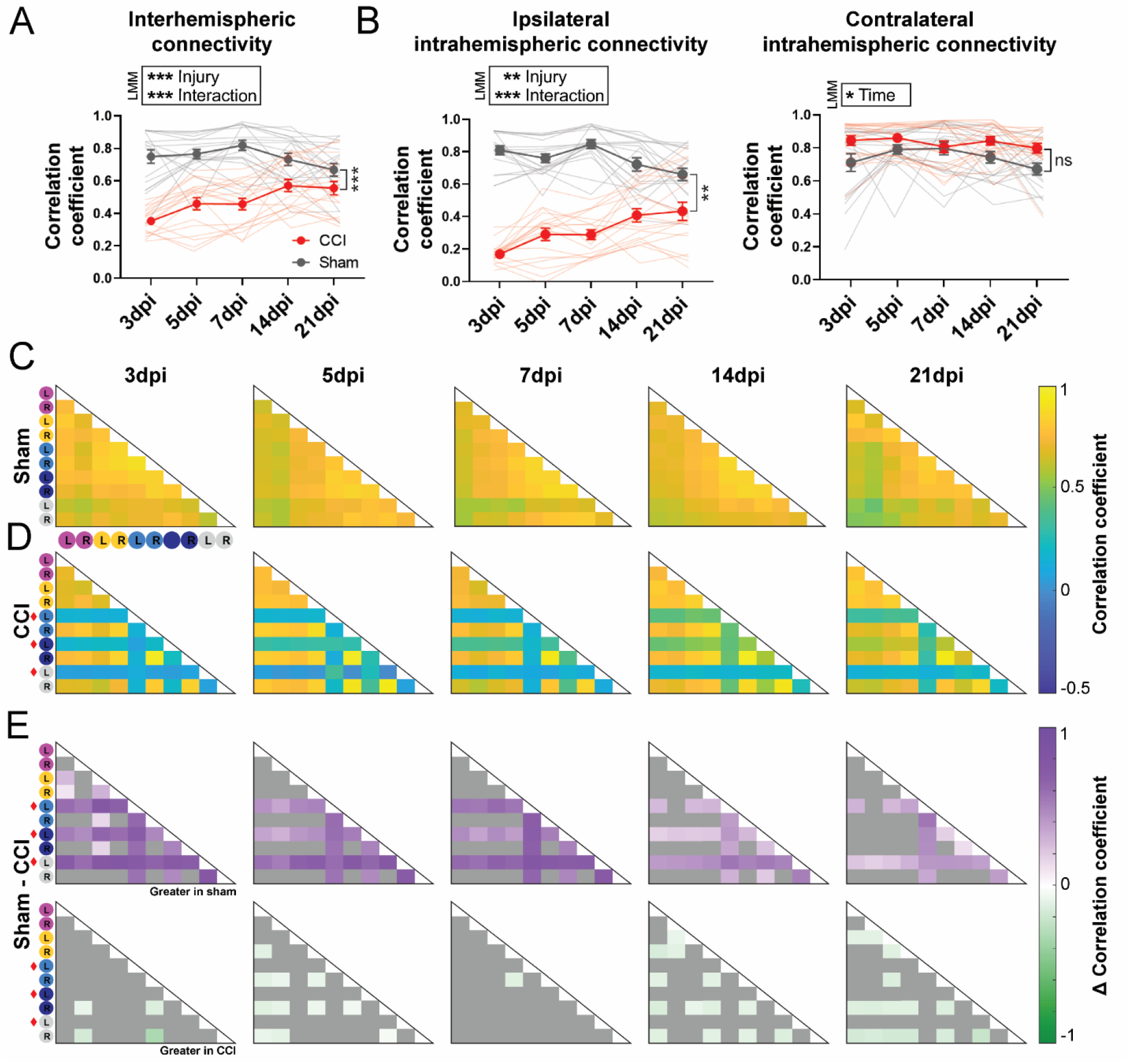
CCI regionallyimpairs functional cortical connectivitythat partiallyrecovers over time. **A.** Interhemispheric connectivity, as calculated by computing the correlation coefficient between anatomic regions in the left versus the right hemisphere. Regions are anterior to posterior: secondary motor cortex, motor barrel cortex, primary somatosensory area forelimb, parietal cortex, and primary visual cortex. Lines indicate each imaging session from all mice contributing to the mean. **B.** Intrahemispheric connectivity in ipsilateral (left) and contralateral (right) hemispheres between the five anatomic regions indicated in A. Connectivity is significantly decreased in the ipsilateral hemisphere but not in the contralateral. For A-B, n = 3 animals per group, 6 trials per animal per timepoint. LMM: *indicates P < 0.05, **indicates P < 0.01, ***indicates P < 0.001. Error bars indicate SEM. **C.** Average connectivity matrices over time in sham and CCI (**D**) mice. This is a point-to-point matrix with each square displaying the correlation coefficient of each anatomic ROI compared to every other ROI. Anatomic ROIs are indicated along the outside of the matrices, marked with an L or R to indicate which hemisphere they originate from, and are anterior to posterior: secondary motor cortex (pink), motor barrel cortex (yellow), primary somatosensory area forelimb (light blue), parietal cortex (dark blue), and primary visual cortex (gray). Areas proximal to CCI are indicated with red diamond. For all conditions in C and D, n = 3 animals per group, 6 imaging trials per timepoint. **E.** Average differences in correlation coefficient between sham and CCI are shown for connections that are significantly decreased (top row) or increased (bottom row) with injury (P < 0.01, Benjamini–Yekutieli corrected Wilcoxon rank sum test (unpaired)). Nonsignificant values are shown in gray. Dpi, days post - injury.

To determine whether specific functional connections are disrupted by CCI, we generated connectivity matrices displaying the functional connectivity between each region and every other region. The connectivity matrices in sham mice showed that the functional connectivity across the brain was stable over all timepoints examined (Figure 5C). Following CCI, the disruption of functional connectivity occurred most notably in the three regions proximal to injury (somatosensory, parietal, and primary visual cortices), but gradually recovered over time for some connections (Figure 5D). These connectivity matrices were consistent both across repeated imaging sessions for individual animals and between animals (Figure S1). We next quantified the difference in functional connectivity between sham and CCI mice over all timepoints and found that all regions proximal to CCI were significantly decreased compared to corresponding regions in shams at 3, 5, and 7 days post-implant (Figure 5E, top row). By 14 days, some regions near CCI were no longer significantly different from shams, specifically in somatosensory and parietal cortices. Interestingly, we found several regions in which functional connectivity was increased in CCI animals, including areas contralateral and anterior to the region of injury (Figure 5E, bottom row).

Given the region-specific recovery of functional connectivity in somatosensory and parietal cortices, we further examined brain areas closest to CCI to determine if these regions displayed unique recovery trajectories over time. We computed the mean functional connectivity between the indicated region proximal to CCI (left somatosensory, parietal, and primary visual cortices) and all other ROIs for each imaging trial (Figure S2). In all three regions, CCI connectivity increased over the 21 days but to different extents with parietal cortex activity most closely approaching sham levels (t_(4)_ = −4.41, p = 9.4e-3 somatosensory; t_(4)_ = −5.1, p = 5.4e-3 parietal; t_(5)_ = −9.25, p = 1.6e-4 visual; effect of injury for all, LMM). This increase in connectivity proximal to CCI further supports our findings that connectivity deficits are driven by focal injury effects. Together, these findings show that functional connectivity is significantly disrupted by CCI, and gradually recovers over time.

### Injury decreases ECoG theta power during stillness but not locomotion

We next examined ECoG data collected during imaging. Due to known state-dependent changes in brain activity, we separated the data into times of locomotion and stillness.^62,63^ Stillness recapitulates aspects of resting state in human fMRI studies^64,65^ and low arousal state in rodent studies.^66^ Movement is considered a higher arousal state and results in distinct circuit changes.^22^ We focused on ECoG theta power (6-8 Hz), which is known to be modulated by state as well as TBI. To control for potential injury-induced changes in animal behavior, we determined the percent time animals spent moving during each imaging session. There was no effect of injury or time post-injury on the total amount of time animals spent moving (t_(11)_ = −0.85, p = 0.45 effect of injury; t_(22)_ = 2.0, p = 0.06 effect of time; t_(22)_ = 0.5, p = 0.62 interaction effect, LMM; Figure 6B).

**Figure 6.**
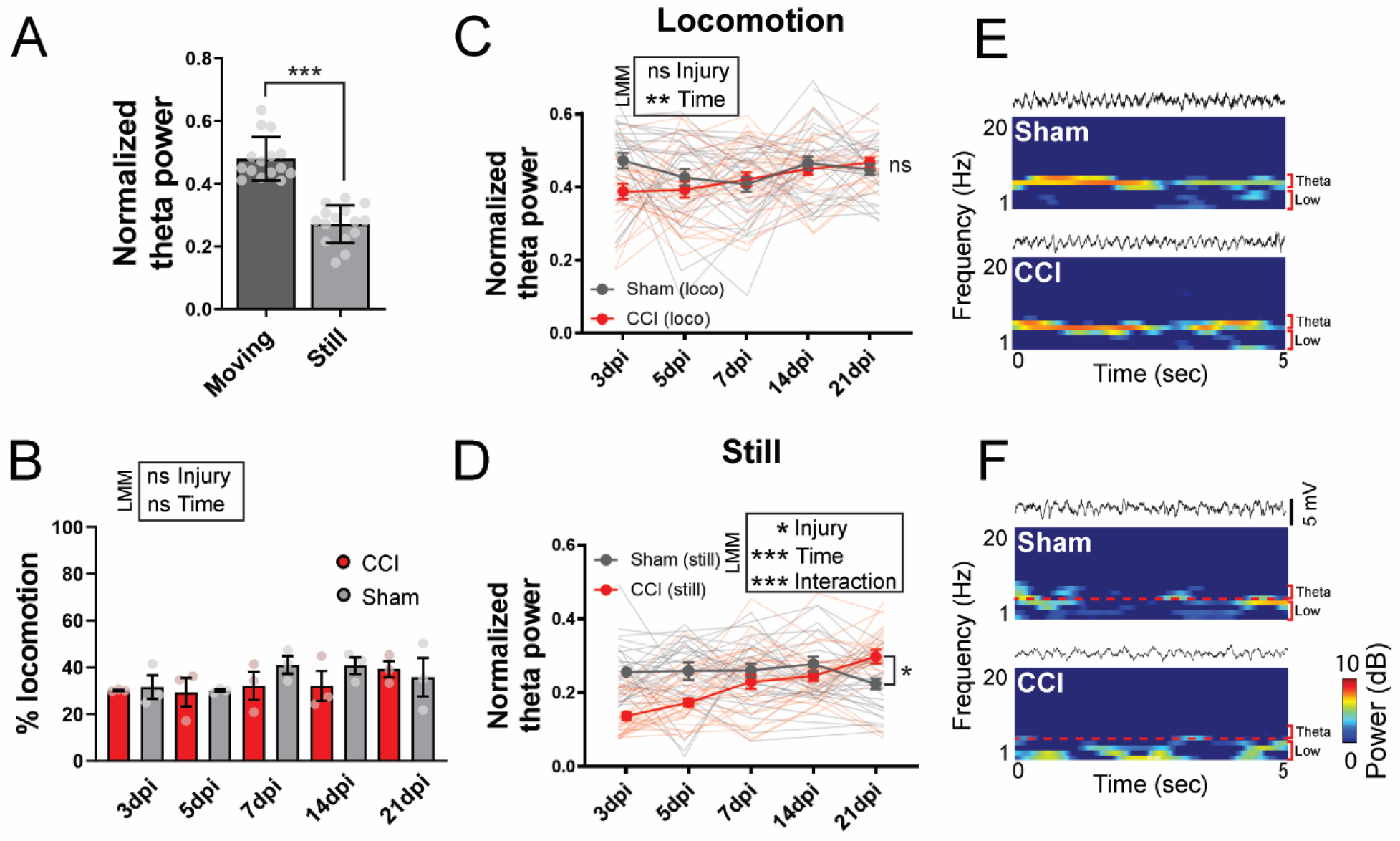
Injury decreases ECoG theta power during stillness but not locomotion. **A.** Quantification of theta power in sham mice from the ipsilateral (left) hemisphere during locomotion and stillness shows a significant decrease in theta power during stillness. Individual points show the average of each mouse at each timepoint. LMM: ***indicates P < 0.001. **B.** Percent time spent moving throughout entire recording session was not significantly different between sham and CCI mice. LMM: ns is not significant. **C.** Theta power during locomotion alone shows no significant differences between CCI and sham mice (ECoG power is average of ipsilateral and contralateral electrodes). **D.** Theta power during stillness alone shows a significant decrease in CCI mice compared to sham. For all conditions, n = 3 animals per group, 3-15 instances of locomotion or stillness at each timepoint. LMM: *indicates P < 0.05, **indicates P < 0.01, ***indicates P < 0.001, ns is not significant. Error bars indicate SEM. **E.** Sample ECoG traces from the ipsilateral (left) somatosensory cortex and corresponding spectrograms for sham (top) and CCI (bottom) mice 3dpi during locomotion show a strong theta band (5-8 Hz). **F.** Sample ECoG traces from the ipsilateral (left) somatosensory cortex and spectrograms for sham (top) and CCI (bottom) mice 3dpi during stillness show lower theta power in CCI mice than in sham. For E and F, theta and low power bands are noted to the right of spectrograms. Loco indicates locomotion. Dpi, days post-injury.

To quantify ECoG during each state, segments of locomotion or stillness were identified, and state transitions removed. In both sham and CCI animals, increased theta power was associated with locomotion (t_(274)_ = 15.6, p < 2e-16, effect of state, LMM; Figure 6A). We next examined thetapower during locomotion and stillness over all timepoints. There was no difference in theta power during locomotion between CCI and sham mice, but theta power did increase slightly over time in CCI mice (t_(11)_ = −1.9, p = 0.08 effect of injury; t_(275)_ = 3.2, p = 1.6e-3 effect of time, LMM; Figure 6C, E). During stillness, however, theta power was significantly decreased in CCI animals compared to shams (t_(5)_ = −3.1, p = 0.03 effect of injury; t_(269)_ = 4.4, p = 1.5e-5 effect of time; t_(269)_ = −4.3, p = 9e-11 interaction effect, LMM; Figure 6D, F). This stillness-specific decrease in theta power in CCI mice returned to normal by 21 days post-injury. In summary, CCI affected ECoG theta power during stillness but not locomotion, suggesting that in vivo studies of brain activity after injury would benefit from state-dependent analysis.

### Locomotion and stillness engage distinct functional connectivity states that are disrupted by CCI

Given our findings of state-dependent and injury-induced changes on ECoG as well as the literature suggesting state-dependent modulation of functional connectivity,^19,22,39^ we quantified changes in functional connectivity during locomotion and stillness in CCI and sham-injured mice. We first determined the interhemispheric correlation coefficients during locomotion and stillness. In sham mice, functional connectivity was significantly higher during stillness than locomotion at all timepoints examined (t_(265)_ = −4.3, p = 2.7e-5 effect of state, t_(265)_ = −0.13, p = 0.9 effect of time, LMM; Figure 7A). In CCI-injured animals, this state-dependent difference in interhemispheric functional connectivity was lost, but connectivity increased for both states over time after injury (t_(271)_ = −2.0, p = 0.051 effect of state, t_(271)_ = 6.1, p = 4.2e-9 effect of time, LMM; Figure 7D).

**Figure 7.**
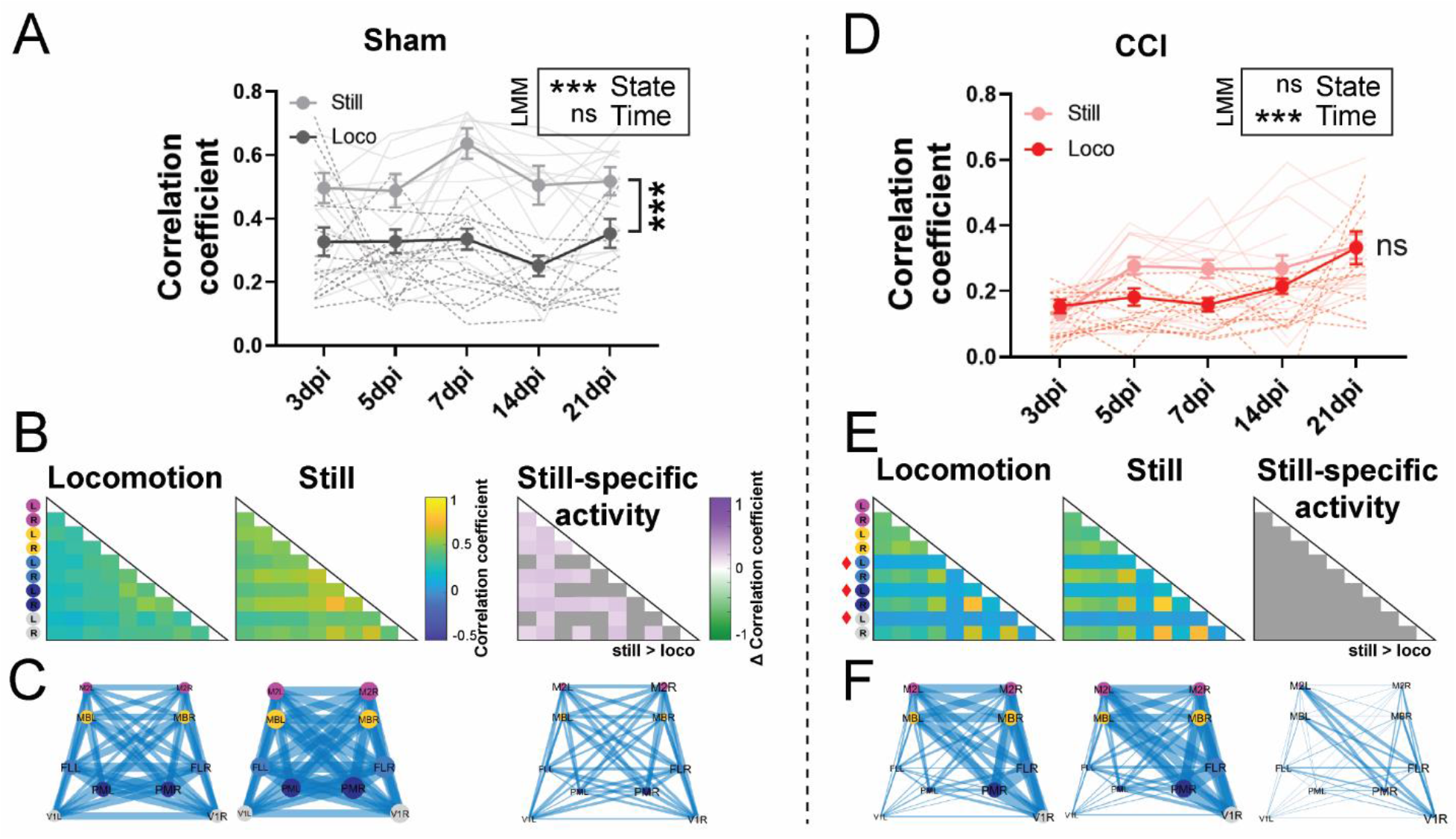
Locomotion and stillness engage distinct functional connectivity states that are disrupted by CCI. **A.** Interhemispheric correlation coefficients derived from mesoscale imaging in sham mice at all timepoints during locomotion and stillness show that stillness engages significantly higher connectivity across all timepoints. LMM: ***indicates P < 0.001, ns is not significant. **B.** Average connectivity matrices (across all timepoints, all trials) in sham mice during locomotion (left) and stillness (middle). Right: Specific connections that are significantly increased during stillness in sham mice (P < 0.01, Benjamini–Yekutieli corrected Wilcoxon signed rank test (paired), n = 3 mice). Nonsignificant values are shown in gray. **C.** Network diagrams indicating the strength of connectivity (the thicker the blue line the stronger the connectivity) in sham mice between the marked anatomic locations during locomotion (left), stillness (middle), and still-specific (still – move) activity (right). **D.** Interhemispheric correlation coefficients in CCI-injured mice at all timepoints during locomotion and stillness. There is no significant difference in connectivity between states in CCI mice. LMM: ***indicates P < 0.001, ns is not significant. **E.** Average connectivity matrices (across all timepoints, all trials) in CCI mice during locomotion (left) and stillness (center). Areas proximal to CCI are indicated with red diamonds. Right: average differences in correlation coefficient between stillness and locomotion in CCI mice (P < 0.01, Benjamini–Yekutieli corrected Wilcoxon signed rank test (paired), n = 3 mice). No connections showed state-dependent changes in CCI mice (gray). **F.** Network diagrams in CCI mice during locomotion (left), stillness (middle), and still-specific (still – move) activity (right). For all conditions, n = 3 animals per group, 3-15 instances of locomotion or stillness at each timepoint. Error bars indicate SEM. Loco indicates locomotion. Dpi, days post-injury.

To examine specific connections engaged in each state-dependent network, we generated separate connectivity matrices for locomotion and stillness. Similar to interhemispheric connectivity, we found a state-dependent change in functional connectivity in sham, but not CCI mice (Figure 7B, E). We then calculated the difference between the average connectivity matrices in locomotion versus stillness to identify a functional signature of brain activity during stillness. In sham mice, there were multiple functional connections particularly between motor and somatosensory regions that showed increased functional connectivity during stillness (Figure 7B). In CCI mice, however, there were no connections that showed increased functional connectivity during stillness (Figure 7E).

We next generated network diagrams of stillness-specific connectivity (Figure 7C, F). The line thickness between brain regions corresponds to the functional connectivity strength. In shams, network diagrams showed uniformly higher connectivity between brain regions during stillness compared to locomotion. After CCI, there was little difference in connectivity between locomotion and stillness. Together, these findings show that in sham-injured mice, functional cortical connectivity is higher during stillness than during locomotion, but in injured mice this state-dependent difference in connectivity is lost at all timepoints examined.

## DISCUSSION

We report a novel approach to perform mesoscale cortical imaging and ECoG after CCI, allowing a previously unattainable view of mouse brain function after TBI. First, we determined that mesoscale implant slightly increased astrocyte reactivity after TBI, but those changes were very small compared to the effect of injury itself. Next, we showed that CCI reduced neuronal activity and functional connectivity in regions proximal to injury. Functional connectivity, but not total neuronal activity, gradually returned over time proximal to injury. This suggests functional reorganization occurs in injured regions even while total neuronal activity remains low. These data are consistent with others’ findings that the amplitude of activity and measurements of connectivity are independent. ^22,24^ Interestingly, total activity and connectivity was increased in CCI mice contralateral to injury, suggesting hyperconnectivity after TBI could contribute to hyper-synchronization associated with post-traumatic epilepsy. These findings suggest that functional connectivity changes dynamically throughout the entire cortex after injury with unique responses in ipsilateral and contralateral hemispheres.

Furthermore, we observed state-specific effects of injury on ECoG activity and functional connectivity, as well as a loss of functional state switching after TBI. Injury decreased ECoG theta power during periods of stillness but had no effect on periods of movement. On imaging in sham mice, functional connectivity was increased at rest and decreased during locomotion, however CCI eliminated this state-dependent change. By 21 days post-injury, CCI connectivity had returned to sham levels during locomotion, but injured animals never regained the state-dependent switch. Together, these findings point to an injury-induced disruption of resting state networks with much smaller effects on locomotion-specific networks, and a loss of state-dependent switching of functional connectivity (Figure 8).

**Figure 8.**
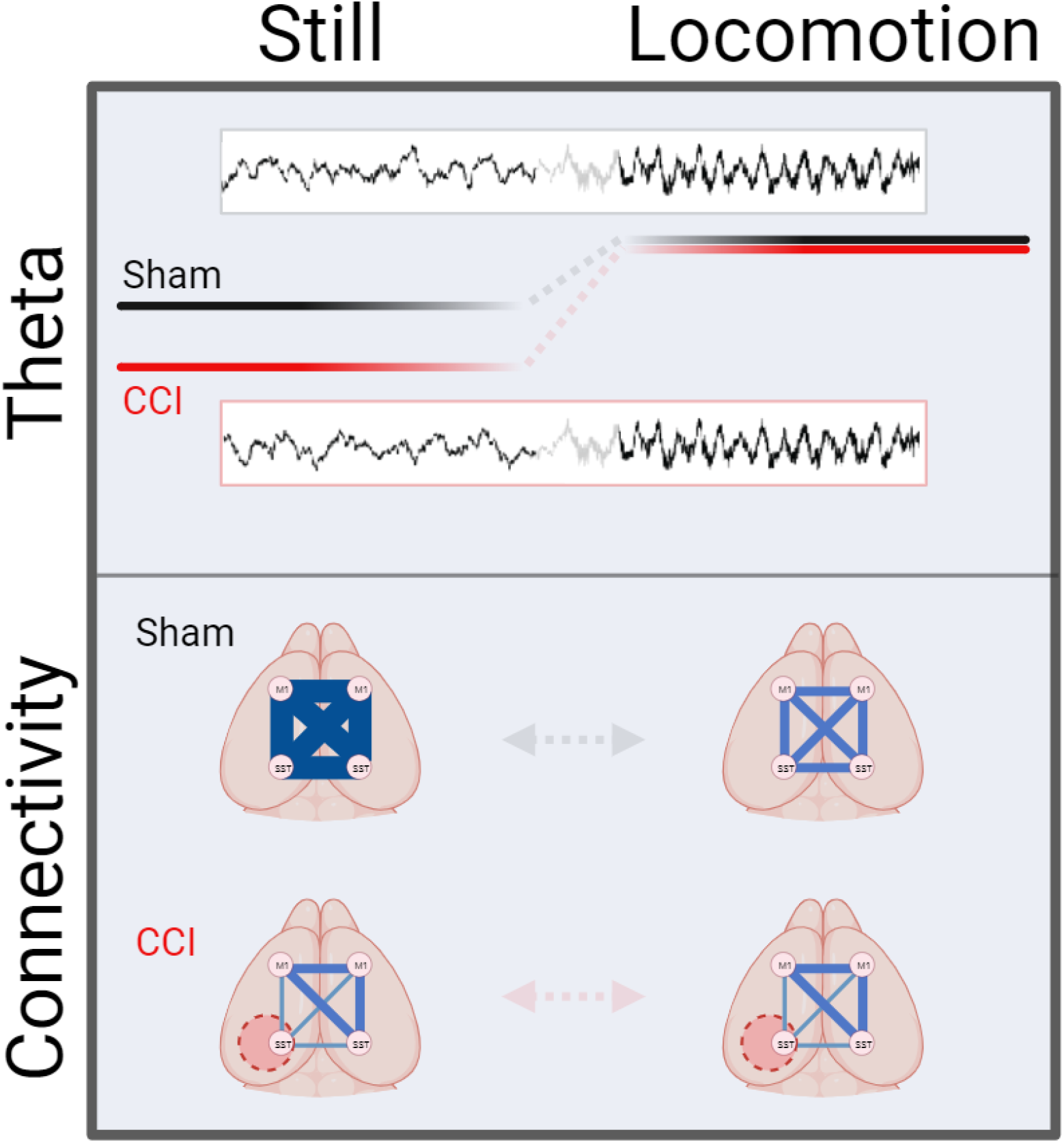
TBI disproportionately alters activity during stillness compared to locomotion. During stillness, CCI decreases theta power and functional connectivity compared to sham levels. During locomotion, theta power is not significantly different from sham levels and functional connectivity is only slightly affected. Theta power and connectivity are altered during stillness but not during locomotion suggesting a state-specific effect of injury.

Our approach permits characterization of a movement state that is challenging to assess using human imaging modalities (e.g. fMRI) and enables study of a large area of cortex after TBI. The connectivity changes we observed in both hemispheres underscore the importance of including regions beyond the perilesional area. Additionally, human studies are limited in that severity, injury region, and etiology differ drastically between patients. Recruiting patients immediately after injury to capture a consistent acute timepoint and following them over time is challenging. fMRI cannot be collected during most motor tasks such as locomotion given the restrictions of the imaging modality. Therefore, our approach uniquely allows quantification of state-specific functional connectivity at multiple timepoints after TBI.

Our findings recapitulate human studies that found arousal state affected both EEG and functional connectivity. EEG and ECoG are modulated by locomotion, particularly in the theta band. Hippocampal theta activity predicts locomotion onset,^62^ and theta dynamics are highly correlated with locomotion speed.^63^ Locomotion-induced theta activity has not been well studied in the cortex. However, cortical neurons are at times phase-locked to hippocampal theta rhythm,^67^ and theta oscillations in the motor cortex coordinate behavior with theta power altered upon locomotion onset.^68^ Our findings build on this, demonstrating in both CCI and sham-injured mice that locomotion corresponds to a significant increase in theta power.

In fMRI studies of healthy participants, functional connectivity decreased with task performance compared to resting state.^39^ During locomotion, human functional connectivity measured via 248-channel EEG was decreased compared to standing particularly in sensorimotor cortex.^19^ This suggests that less coordinated input from the cortex is required for locomotion than for standing. State also alters functional connectivity in mice. Widefield cortical imaging demonstrated globally decreased connectivity between nearly all nodes examined during locomotion compared to rest.^22,40^ Together, this indicates that both movement and task-dependent variations in state alter cortical activity in humans and in mice. Moving forward, it will be important to collect data during multiple states to gain a more complete picture of cortical network activity and how it is modulated by disease. For example, theta relative power has been proposed as a biomarker of epileptogenesis following TBI.^69^ Performing state-dependent analysis of theta power could improve the strength and accuracy of this biomarker.

In addition to state, EEG and functional connectivity are altered with TBI. In rodent models of TBI, theta power^70–72^ and functional connectivity are decreased after TBI.^14,25,26^ In patients with lesions, connectivity was decreased proximal to the lesion compared to intact contralateral regions and controls.^14^ Functional connectivity was also predictive of recovery with patients who exhibited stronger interhemispheric functional connectivity performing better on cognitive tests.^26^ Rehabilitated TBI patients had connectivity profiles more similar to healthy controls than to the non-rehabilitated group and scored higher on cognitive and memory indexes. ^15^ These human and rodent studies indicate that functional connectivity may be an important measure to track disease severity, recovery, and response to rehabilitation. We found dynamic changes in ECoG power and functional connectivity particularly in the first week after injury, suggesting monitoring patients over several timepoints could provide additional insight into their recovery.

Beyond the commonly observed decrease in functional connectivity perilesionally, several rodent studies also observed focal increases in functional connectivity after TBI. This consistently occurred contralateral to injury in tandem with decreased connectivity ipsilateral to injury. ^73,74^ Increased connectivity was observed in humans after mild TBI both between regions in existing brain networks and recruitment of regions not previously in-network, both thought to represent compensation after injury.^75^ This could explain why we observed both increased total neuronal activity and connectivity contralateral to injury. Moving forward, it will be important to understand if contralateral connectivity has implications for functional recovery.

Our study has several caveats. Our imaging approach is restricted to upper-layer cortical activity, and the model of CCI we employ damages the cortex, ^27^ limiting our ability to directly quantify activity in the site of injury. We therefore selected ROIs not directly within the injury site but on the border in the perilesional area. Additionally, GCaMP7c expression is not altered by CCI perilesionally, allowing rigorous quantification of neuronal activity even after injury. Addressing the caveats of imaging neuronal activity after CCI is important because CCI models the sequelae of the most severe TBI patients with the highest symptom burden. ^34,35^ Additionally, during the development of our combined CCI/mesoscale imaging approach, we focused only on male mice. It will be important to include female mice in future studies as TBI occurs in both sexes,^76^ and male and female mice differ in their response to TBI with sex-specific hormones potentially acting neuroprotectively in females.^77^ It will also be important to implement a behavioral assessment of functional recovery that can be evaluated in tandem with functional connectivity. Employing behavioral tests alongside imaging and ECoG in male and female mice will allow us to begin testing functional effects and efficacy of therapeutic interventions. Additionally, further separating stillness epochs into low and moderate arousal states using pupil or facial motion could allow us to detect potential differences in how injured and sham animals engage in stillness. Finally, because this initial study was designed to validate mesoscale imaging in the injured brain, n-values are somewhat low (n = 3 male mice for all assays). Both statistical and power analysis, however, show that our study is sufficiently powered to detect changes in functional connectivity and ECoG after CCI. More subtle changes, with smaller effect sizes, may not be detected in this study and future studies with this now validated approach will include larger sample sizes and interventional approaches.

The most significant and novel finding in our study is demonstrating that both state and TBI alter functional cortical activity and that TBI has much greater effects on networks engaged during stillness as compared to during movement. In addition, we report dynamic recovery of connectivity after TBI driven by the ipsilateral hemisphere, along with increases in connectivity over sham levels in the contralateral hemisphere. Few previous studies have performed state-dependent analysis after TBI. A porcine model of pediatric TBI used fMRI to compare resting and stimulation-induced functional connectivity under anesthesia after injury.^78^ Functional connectivity was decreased in TBI piglets compared to shams and state affected which networks were most decreased after injury.^78^ While anesthesia drastically alters brain activity, this study nonetheless emphasizes that state modulates the location and extent of brain regions affected by TBI. In summary, we report one of the first studies to perform awake state-dependent analysis of TBI-injured animals and identified state-specific changes in connectivity that were altered by CCI particularly when animals were at rest. Future studies will utilize mesoscale cortical imaging to help identify disease-relevant biomarkers that will allow for early and individualized therapeutic interventions.

## TRANSPARENCY, RIGOR, AND REPRODUCIBILITY

Animals were randomized to experimental groups and all CCI and sham surgeries were performed when animals were 8-10 weeks of age. Sample sizes ensure power >0.8 for α=0.05. ROI mapping and identification of periods of stillness and locomotion were automated to avoid experimenter bias. We used linear mixed modeling to account for inter - and intra-animal variability. Detailed statistical analyses are reported in Tables S1 and S2.

## ACKNOWLEDGMENTS

We are grateful to Christopher Shen for his generous help with developing software for locomotion detection.

## AUTHORS’ CONTRIBUTIONS

SBT: conceptualization, methodology, software, investigation, writing - original draft. SC: methodology, validation. JM, MS, LB, TM, SH, MS: resources, writing – review & editing. MW: writing – review & editing. SQ: software, writing – review & editing. FN: formal analysis, writing – review & editing. RO, MJH, JAC: software, resources, writing – review & editing. MA, CD: conceptualization, methodology, software, writing - original draft.

---

**Supplemental Figure 1.**
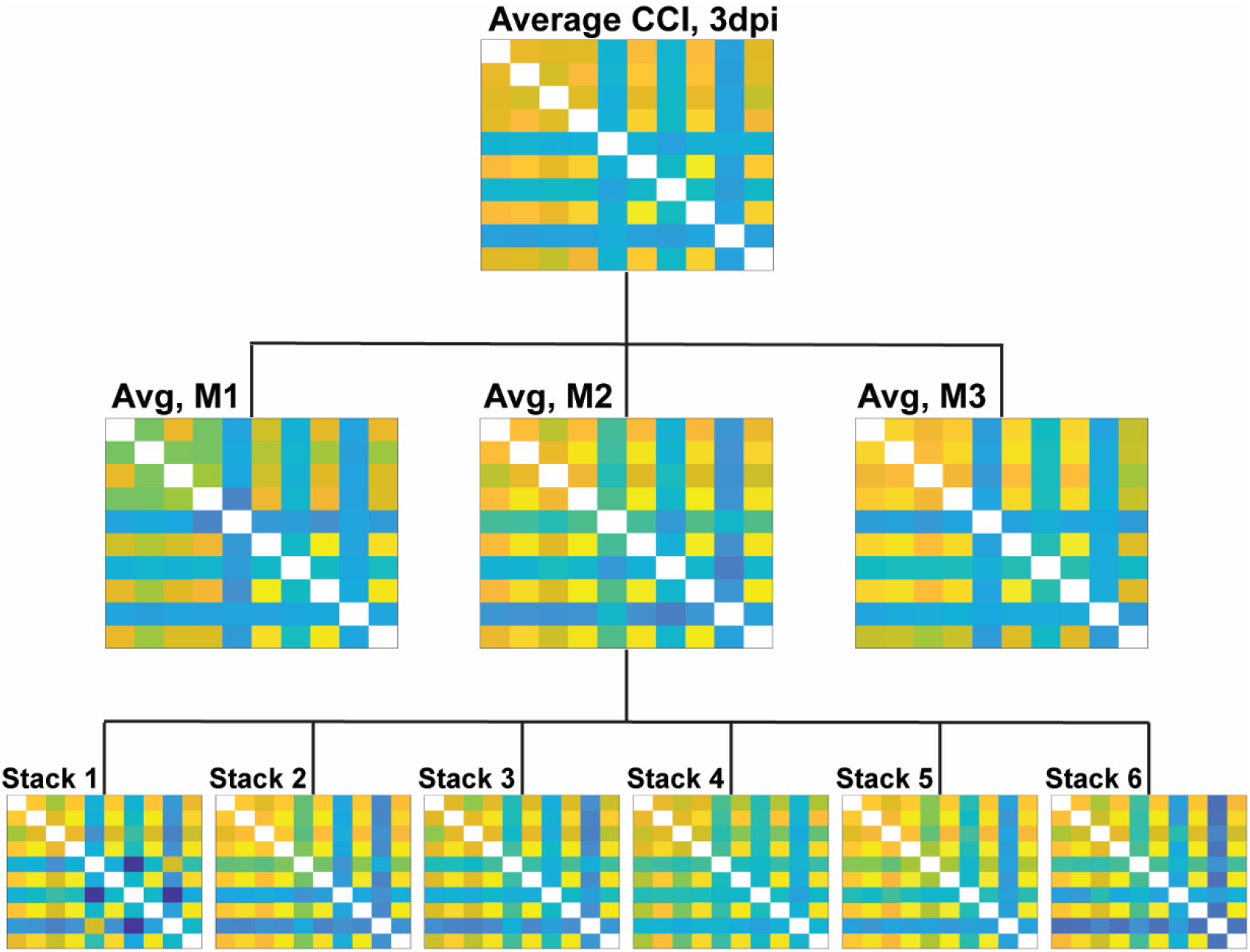
Connectivity matrices have minimal inter- and intra-animal variability. From top to bottom, the average connectivity matrix for CCI animals 3 DPI. Middle row: The average connectivity matrix of each individual mouse (M1, M2, and M3). Bottom row: The connectivity matrices for mouse #2 (M2) for each 75 second imaging session. Dpi, days post-imaging.

**Supplemental Figure 2.**
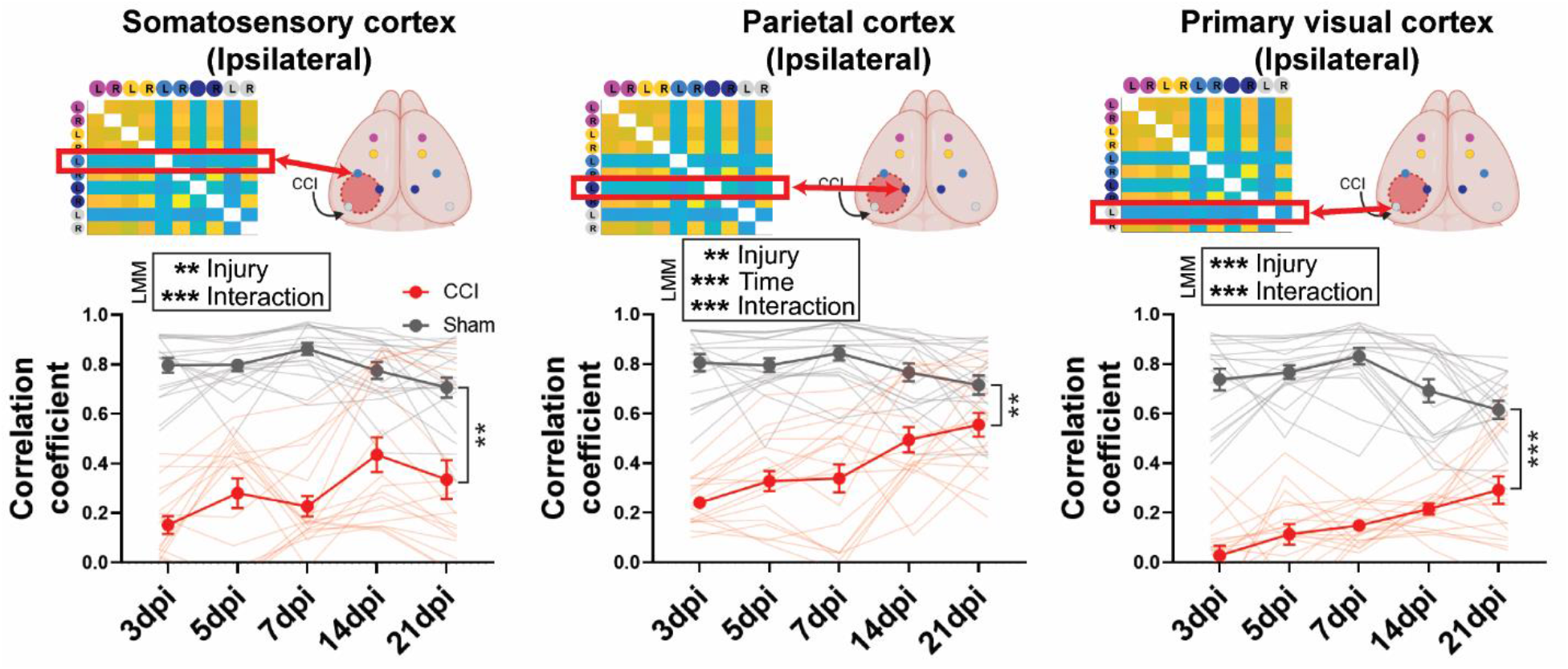
Functional connectivity is decreased perilesionally. Average correlation coefficient from a single ROI compared to all others over time proximal to injury in CCI mice compared to identical ROIs in shams. In all three regions, connectivity is decreased in injured mice. LMM: *indicates P < 0.05, **indicates P < 0. 01, ***indicates P < 0.001. Error bars indicate SEM. Dpi, days post-injury.

**Supplemental Table 1.**
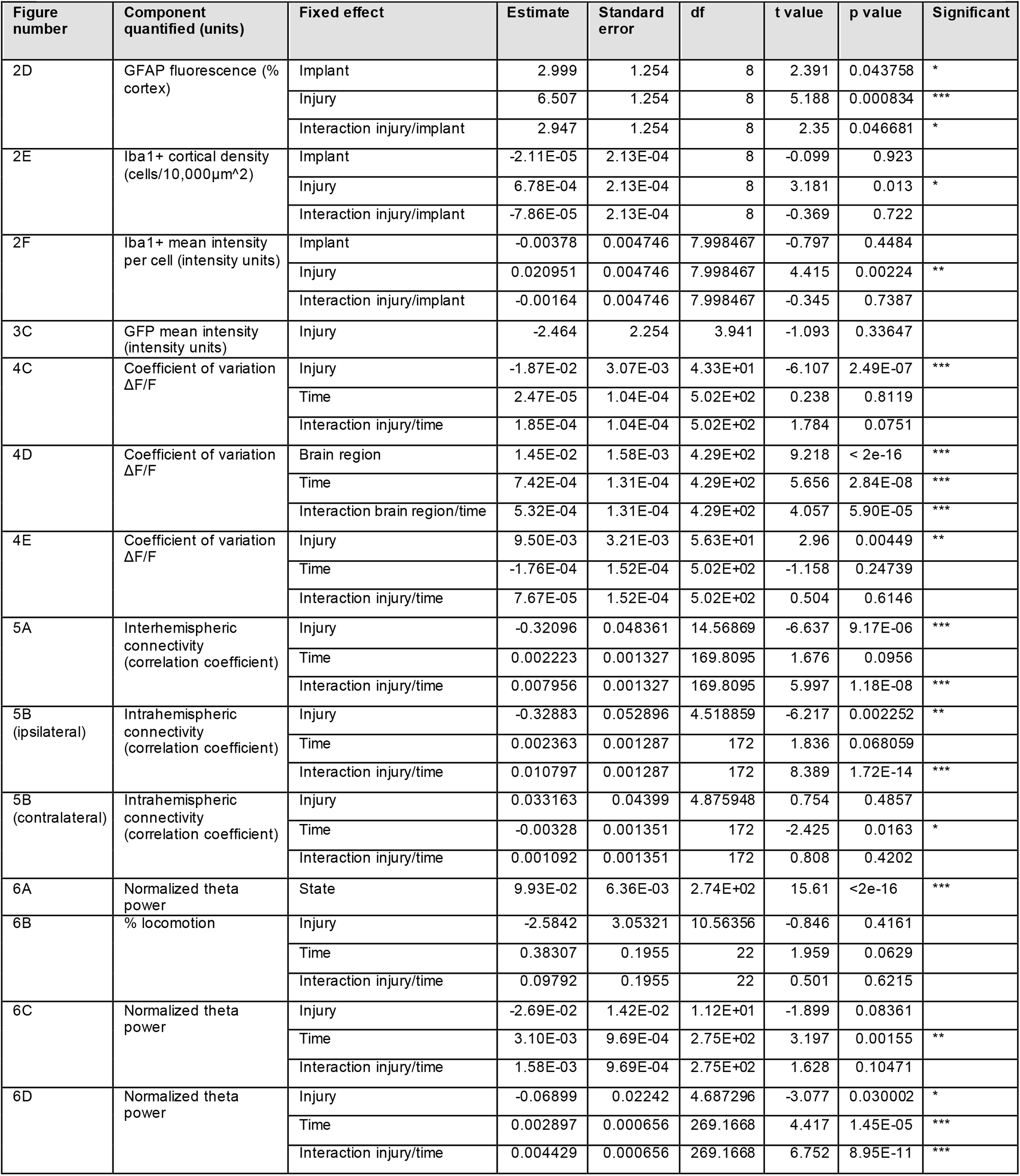
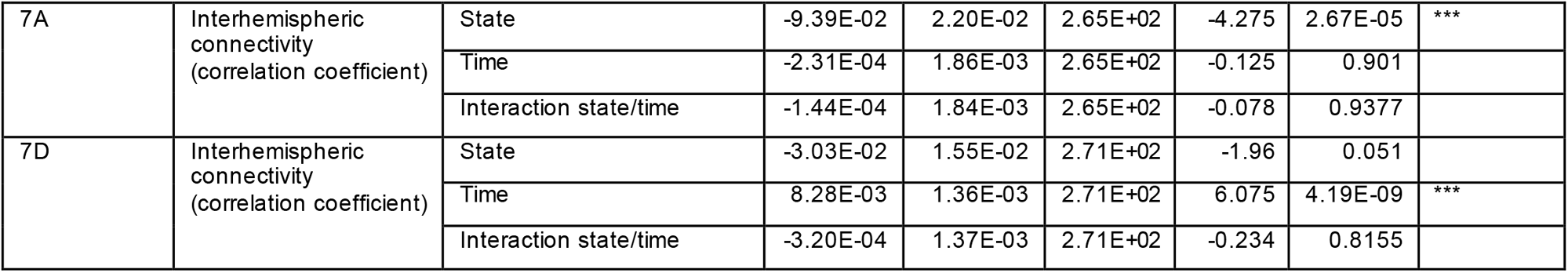
Linear mixed modeling results for figures 2-7, organized by figure.

**Supplemental Table 2.** Linear mixed modeling results for Figure S2.

| Figure number | Component quantified (units) | Fixed effect | Estimate | Standard error | df | t value | p value | Significant |
| --- | --- | --- | --- | --- | --- | --- | --- | --- |
| Supplemental 2 (somatosensory cortex) | Interhemispheric connectivity (correlation coefficient) | Injury | -0.33196 | 0.075257 | 4.383589 | -4.411 | 0.00942 | ** |
|  |  | Time | 0.002292 | 0.001593 | 172 | 1.439 | 0.15189 |  |
|  |  | Interaction injury/time | 0.008093 | 0.001593 | 172 | 5.082 | 9.69E-07 | *** |
| Supplemental 2 (parietal cortex) | Interhemispheric connectivity (correlation coefficient) | Injury | -0.3092 | 0.060756 | 4.412544 | -5.089 | 0.005393 | ** |
|  |  | Time | 0.005756 | 0.00133 | 172 | 4.328 | 2.55E-05 | *** |
|  |  | Interaction injury/time | 0.011241 | 0.00133 | 172 | 8.452 | 1.18E-14 | *** |
| Supplemental 2 (primary visual cortex) | Interhemispheric connectivity (correlation coefficient) | Injury | -0.39414 | 0.042605 | 5.417946 | -9.251 | 0.000161 | *** |
|  |  | Time | 0.002068 | 0.0016 | 172 | 1.292 | 0.197949 |  |
|  |  | Interaction injury/time | 0.010963 | 0.0016 | 172 | 6.852 | 1.25E-10 | *** |

